# Anxiety enhances pain in a model of osteoarthritis and is associated with altered endogenous opioid function and reduced opioid analgesia

**DOI:** 10.1101/2020.04.24.057570

**Authors:** Amanda Lillywhite, Stephen G. Woodhams, David J. G. Watson, Li Li, James J. Burston, Peter R. W. Gowler, Meritxell Canals, David A. Walsh, Gareth J. Hathway, Victoria Chapman

## Abstract

Chronic pain states such as osteoarthritis (OA) are often associated with negative affect, including anxiety and depression. This is, in turn, associated with greater opioid analgesic use, potentially contributing to current and future opioid crises. We utilise an animal model to investigate the neurobiological mechanisms underlying increased opioid use associated with high anxiety and chronic pain.

Combining a genetic model of negative affect, the Wistar Kyoto (WKY) rat, and intra-articular injection of monosodium iodoacetate (MIA; 1mg), our model of high anxiety and augmented OA-like pain behaviour mirrors the clinical problem. Effects of morphine (0.5-6mg.kg^-1^) on pain behaviour and spinal nociceptive neuronal activity were determined in WKY rats, and normo-anxiety Wistar rats, 3 weeks after MIA injection. WKY rats developed augmented OA-like pain, and had blunted inhibitory responses to morphine, when compared to Wistar rats. Potential alterations in endogenous opioid function were probed via systemic blockade of opioid receptors with naloxone (0.1-1mg.kg^-1^), quantification of circulating levels of β-endorphin, and determination of spinal expression of the mu-opioid receptor (MOR). These studies revealed increased opioidergic tone, and increased spinal desensitization of MORs via the master phosphorylation site at serine residue 375, in this model.

We demonstrate attenuated MOR function in the absence of previous exogenous opioid ligand exposure in our model of high anxiety and OA-like pain, which may account for reduced analgesic effect of morphine and provide a potential explanation for increased opioid analgesic intake in high anxiety chronic pain patients.

**Significance Statement:** Chronic pain affects large numbers of people, and pain management often relies on poorly effective opioid analgesics, the iatrogenic effects of which are increasingly recognised. The endogenous opioid system - the target for exogenous opioid analgesics - plays key roles in emotional affective states and pain control, but the complex interplay between anxiety, chronic pain, and endogenous opioid system function is challenging to study in people. Here, we have addressed this using a clinically-relevant experimental model. Anxiety-like behaviour was associated with increased chronic arthritis-like pain behaviour, altered opioid receptor function, and reduced efficacy of opioid analgesics. We provide new evidence, which may explain why chronic pain patients with comorbid high anxiety have higher opioid analgesic use.

## 1. Introduction

Opioid analgesics have formed the backbone of the management of moderate to severe pain for decades. Opioid drugs, such as morphine, produce their effects via μ-opioid receptors (MOR) at key sites in the spinal cord and brain (McMahon, 2013). Since the 1990s, there has been a shift in the prescribing of opioids from managing acute pain and pain in terminally ill patients, to more wide-spread prescribing for a wide range of long-term pain conditions (Curtis et al., 2019).

Both morphine’s analgesia and its side-effects (tolerance, respiratory depression, euphoria, and dependence) are mediated by MOR. Clinical studies provide strong evidence that, although opioids are excellent analgesics for acute pain such as post-operative pain, long-term use of opioids is not associated with useful pain relief in most people (Colvin et al., 2019). This loss of analgesic benefit is attributed to MOR tolerance and the manifestation of opioid-induced hyperalgesia (Yang et al., 2019), both of which have major implications for pain trajectories *per se*, as well as postoperative outcomes and a contribution towards poorly-controlled pain (Colvin et al., 2019). Chronic pain alters endogenous opioid function, including increased release of endogenous opioid ligands, such as β-endorphin, which may lead to alterations at MOR with increased phosphorylation, opioid tolerance, and lower analgesic responsiveness to morphine (Petraschka et al., 2007; Bruehl et al., 2013).

Osteoarthritis (OA) is the most common form of arthritis (Glyn-Jones et al., 2015) and the fastest growing cause of chronic pain worldwide (Vos et al., 2012; GBD_2013_DALYs_&_HALE_Collaborators et al., 2015). OA pain phenotypes reflect both nociceptive and neuropathic mechanisms of pain (Glyn-Jones et al., 2015; Perrot, 2015), and approximately 20% of people exhibit features of central sensitization, including regional spread of pain (Suokas et al., 2012; Arendt-Nielsen et al., 2015). Despite the reportedly high rates of opioid-prescribing for OA pain (Bedson et al., 2016; Thorlund et al., 2019), opioids have no superior effect over non-opioid treatments over 12 months (Krebs et al., 2018), and long-term opioid use is associated with increased risk of adverse events (Bedson et al., 2016). Negative affect, including anxiety and depression, is associated with exacerbated chronic pain (de Heer et al., 2014), and is common in people with OA (Axford et al., 2010; Barnett et al., 2018). Previous studies reported complex relationships between endogenous opioid function and depressive symptoms and trait anxiety in people (Burns et al., 2017), and negative affect is associated with greater use of opioid analgesics in people with OA (Valdes et al., 2015; Barnett et al., 2018; Namba et al., 2018). Substantial advances in the understanding of the mechanisms of pain and opioid-induced analgesia have been made using rodent models of disease and molecular and pharmacological studies of opioid receptor function (Pasternak, 2012, 2014). Inbred Wistar Kyoto (WKY) rats are used as an experimental model of anxiety-like behaviour (McAuley et al., 2009). Previously, we reported exacerbated pain behaviour in the monosodium iodoacetate (MIA) model of OA pain in WKY rats, including a spread of pain to the contralateral side and increased markers of astrocytic activation (Burston et al., 2019).

We hypothesised that heightened anxiety is associated with alterations in the endogenous opioidergic system and MOR function, which may counter the effects of exogenous morphine used to treat OA pain. Herein, we evaluated the degree of morphine-mediated analgesia and effects of naloxone-mediated blockade of the endogenous opioid system in a model of OA-like pain in high anxiety WKY and normo-anxiety Wistar rats, then investigated the underlying neurobiological mechanisms. Circulating levels of β-endorphin were measured in Wistar and WKY rats, and potential changes in MOR function in the spinal cord were investigated. Levels of MOR protein and MOR receptor phosphorylation at serine residue 375, which is required for morphine-mediated desensitization (Schulz et al., 2004), were quantified in the dorsal horn of the spinal cord.

## 2. Materials & Methods

### 2.1 Experimental Animals

Studies were in accordance with UK Home Office Animals (Scientific Procedures) Act (1986) and ARRIVE guidelines (Kilkenny et al., 2012). A total of 183 male rats were used for this study; Wistar *n* = 97 (Charles River, Margate, United Kingdom), & Wistar Kyoto *n* = 86 (WKY; Envigo, Bicester, United Kingdom). Males only were used to reduce variability in the data, and to maintain consistency with previous studies characterising this model. Wistar rats were used as the most genetically similar control strain to WKY. Rats were group housed by strain in groups of 4 on a 12-hour light/dark cycle in a specific pathogen-free environment with *ad libitum* access to standard rat chow and water. Treatments were randomly assigned to rats, and experimenters were blinded to all treatment groups throughout the study. A total of 14 rats were excluded from the study (7.6%; see **Extended Data Table 1-1** for further details).

### 2.2 Induction of the MIA model of OA pain

All rats received a single intra-articular injection into the left knee, through the infra-patellar ligament using a 29-gauge hypodermic needle, under isoflurane anaesthesia (3% ILmin^-1^ O2). Rats were randomly assigned to receive either 1mg/50μl MIA in 0.9% saline (Wistar *n* = 48, WKY *n* = 45) or 50μl 0.9% saline (Wistar *n* = 41; WKY *n* = 41). Health and welfare checks were performed immediately after anaesthetic recovery, then daily for the first 3 days, and weekly thereafter. Pain behaviour was assessed twice weekly from D3 to 21 post-model induction.

### 2.3 Behavioural Testing

All rats were habituated to the pain behaviour testing environment for 2 days prior to taking baseline measurements to minimise any exploratory behaviour during testing. Baseline measurements were taken in the morning prior to treatment (D0). Weightbearing (WB) asymmetry was assessed using an incapacitance tester (Bove et al., 2003) (Linton Instrumentation, Diss, UK). Healthy rats distribute their weight evenly between limbs, and a weight shift onto the contralateral limb indicates pain at rest in the ipsilateral knee joint (Bove et al., 2003). Referred pain at distal sites was assessed via determination of mechanical hindpaw withdrawal thresholds (PWTs) via von Frey hair (vFH) monofilaments using the up/down method for both ipsilateral and contralateral paws (Chaplan et al., 1994).

The elevated plus maze (EPM; Walf and Frye, 2007) was used to measure anxiety in Wistar and WKY rats. Rats were placed into the centre of the arena with their nose pointing into an open arm and activity was monitored for a period of 10 minutes using Ethovision software (Noldus Information Technology, Netherlands). Latency to enter the open outer arm and total time spent in closed versus open arms was then determined. Some exploratory behaviour in the open arms of the maze is expected in normo-anxiety animals, whilst restriction of activity to the closed arms is considered a surrogate indicator of anxiety-like behaviour.

To control for any strain differences in locomotor activity in a non-anxiogenic environment, the number of beam breaks per hour was assessed in an activity box (39.5cm x 23.5cm x 24.5cm, 4 x 8 photobeam array; Photobeam Activity System, San Diego Instruments, USA) in animals of both strains at baseline and 19-21 days after model induction (Pezze et al., 2014).

### 2.4 Pharmacological Interventions

#### 2.4.1 Systemic naloxone/morphine behavioural study

To assess differences in sensitivity to systemic opioids, and potential alterations in endogenous opioidergic tone, behavioural responses to cumulative systemic doses of morphine, naloxone, or vehicle (0.9% saline) were determined in Wistar and WKY rats 21 days after model induction in separate groups of rats, respectively. Pre-drug pain behaviour was assessed via WB and vFH in MIA or saline-treated rats, then following 3 cumulative systemic doses of morphine (0.5, 2, & 3.5 mg.kg^-1^mL^-1^, s.c.; Wistar/MIA *n* = 10, WKY/MIA *n* = 10) or vehicle (Wistar/saline *n* = 8, WKY/saline *n* = 8), or naloxone (0.1, 0.3, & 1 mg.kg^-1^mL^-1^, s.c.; Wistar/saline *n* = 12, Wistar/MIA *n* = 12, WKY/saline *n* = 10, WKY/MIA *n* = 11). Pain behaviour was assessed at 15, 30, and 60 mins after each dose.

#### 2.4.2 In vivo *Spinal Electrophysiology*

To assess potential differences in spinal sensory network activity and responses to systemic morphine between strains, in the presence and absence of OA-like pain, single unit extracellular recordings were obtained from wide dynamic range (WDR) neurons in the deep dorsal horn, as previously described (Urch and Dickenson, 2003). Briefly, a laminectomy was performed under isoflurane anaesthesia (surgery: 3%, maintenance: 1.5%) to expose lumbar L4 6 spinal cord, and a WDR neurone was located with a receptive field in the toes of the ipsilateral hindpaw via a glass-coated tungsten microelectrode (Wistar/saline *n* = 10, Wistar/MIA *n* = 15; WKY/saline *n* = 12, WKY/MIA *n* = 12). Once identified, WDR neurones were characterised via electrical stimuli delivered to the receptive field via bipolar electrodes. WDRs exhibit responses to electrical stimulation at Aβ, Aδ, and C fibre latencies, and wind up in response to a repeated noxious electrical stimulation (16 x 50ms, 0.5Hz, delivered at 3-fold C fibre threshold). The degree of wind up can be used as a proxy of central sensitization (D’Mello and Dickenson, 2008). Mechanical responses to hindpaw stimulation with 8, 10, 15, and 26g vFH (10s application, 10s inter stimulus interval) were recorded at baseline, and then every 10 mins following cumulative systemic doses of morphine sulfate (0.5, 2.5, & 6 mg.kg^-1^, s.c., 60 min intervals).

### 2.5 Assessment of Opioid Function in *Ex Vivo* Tissues

#### 2.5.1 Western Blotting

To determine potential differences in spinal opioid receptor expression, fresh spinal cord tissue was collected from WKY and Wistar rats 21 days after intra-articular injection of MIA or saline (*n* = 4/strain). Rats were killed via overdose with sodium pentobarbital (Euthatal, 2mL, i.p.), decapitated, and spinal cord tissue rapidly collected via hydraulic extrusion. The lumbar enlargement of the spinal cord was then hemisected down the midline, snap-frozen in liquid nitrogen, and stored at −80°C until processed. Tissue was homogenised in RIPA buffer with added protease and phosSTOP inhibitor cocktails (Sigma Aldrich, Gillingham, UK) to prevent degradation of proteins and preserve their phosphorylation sites. An equal amount of protein (150μg) from each sample was then separated via SDS-PAGE, transferred onto nitrocellulose membranes, and probed for expression of total MOR (rabbit anti-mu opioid receptor, Neuromics, RA10104, 1:500), P-ser375 MOR (rabbit anti-mu opioid receptor Ser375, BIOSS-Stratech, bs-3724R, 1:500), and β-actin (mouse anti-β-actin, Sigma, A5441, 1:5000) via overnight incubation in 5% milk at 4°C. Secondary antibodies were IRDye donkey anti-rabbit 800CW and donkey anti-mouse 680RD (1:5000 in 5% milk, RT, 1.5hr), and resulting fluorescent signal imaged via Licor Odyssey system (LI-COR Biosciences, Cambridge, UK) and resulting bands quantified via densitometry measurements in Image Studio Lite version 5.2 (LI-COR Biosciences). Data are expressed as expression level relative to β-actin.

#### 2.5.2 β-endorphin ELISA

To determine any strain differences in circulating endogenous opioid peptides, levels of β-endorphin were measured via a commercial ELISA in blood obtained from naïve male Wistar (*n* = 8) and WKY (*n* = 7) rats. Rats were humanely killed via overdose of sodium pentobarbital (Euthatal, 2mL, i.p.) and exsanguinated to generate blood samples. Whole blood was collected in ice cold tubes containing 200μl heparin and processed within one hour. Samples were centrifuged at 13,000 rpm, 4°C for 20 mins, and the supernatant plasma collected and stored at −80°C prior to assay. Plasma samples were assayed for β-endorphin in duplicate using a commercially available EIA kit (Phoenix Pharmaceuticals, Burlingame, CA, USA) according to the manufacturer’s instructions.

### 2.6 Assessment of joint pathology

At the end of each study, rats were euthanised via overdose with sodium pentobarbital (Euthatal, 2ml, i.p.) and whole knee joints collected and fixed in 10% neutral buffered formalin (Guingamp et al., 1997). Knee joints were disarticulated, and cartilage damage quantified by a blinded experimenter via established macroscopic scoring methods (Guingamp et al., 1997). 7.6% of rats were excluded from the study due to inconsistent joint pathology (see **Extended Data Table 1-1** for further details).

### 2.7 Experimental design & statistical analyses

#### 2.7.1 Behavioural Data

To correct for strain differences in total bodyweight, weightbearing asymmetry was calculated as the percentage of weight borne on the ipsilateral hindlimb compared to the total of both hindlimbs [ipsilateral g/(ipsilateral g+contralateral g)]. Similarly, locomotor activity was assessed as the number of beam breaks per minute per kilogram bodyweight. Mechanical PWTs are reported as raw vFH values, or as change in the number of vFHs from baseline when pooling between different cohorts of animals (see Figure 1). For response to pharmacological interventions, % analgesia was calculated for both weightbearing and PWTs. 100% analgesia constitutes total normalisation of weightbearing asymmetry [ipsilateral g/(ipsilateral g+contralateral g) = 50%] or a normalisation of PWT to pre-model baseline values. Anxiety-like behaviour was determined for each animal via Ethovision software as the area under the curve (AUC) of the time spent in the open arms of the elevated plus maze in 1 minute bins for the entire 10 min assessment period, and latency to enter the open outer arm in seconds.

**Figure 1.**
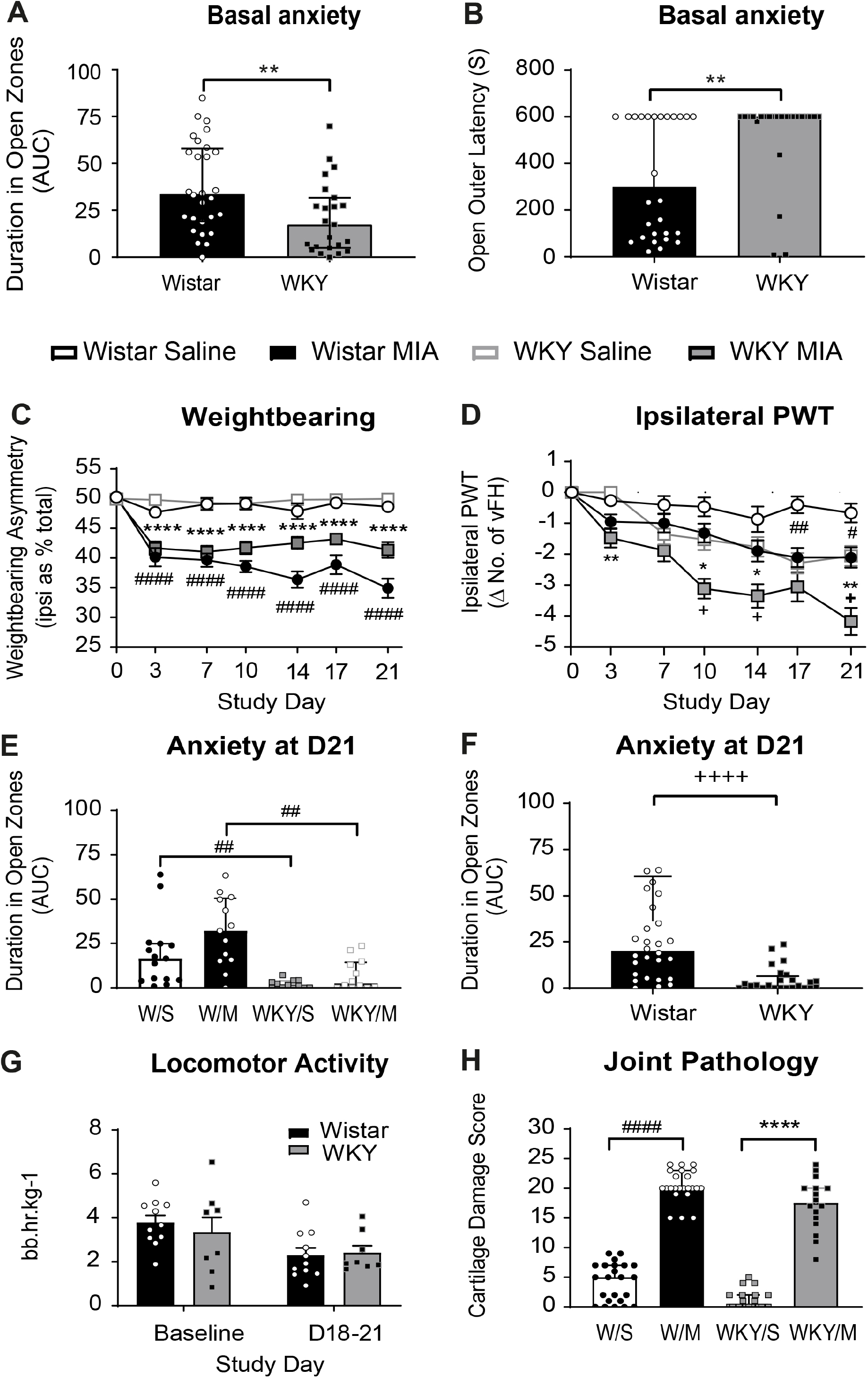
Anxiety-like phenotype & exacerbated OA-like pain in the WKY-MIA model. Basal anxiety-like phenotype in WKY rats in the elevated plus maze (EPM). WKY rats spent significantly less time in the open arms of the maze (**A**), and showed an increased latency to enter the most anxiogenic area of the arena; the open outer arms (**B**). Bars indicate median values, error bars represent IQR. ** p<0.01 versus Wistar rats, one-tailed Mann-Whitney *U* test. MIA-treated rats had a similar degree of weight-bearing asymmetry in both the Wistar and WKY strains (**C**), with pain behaviour evident from day 3 onwards and maintained until post-injection day 21 (D21). Saline administration did not effect weightbearing in either strain. Data are mean ± SEM % weight borne on the ipsilateral hindlimb, #### p< 0.001 versus Wistar saline, **** p< 0.0001 versus WKY saline, 2 way ANOVA with Tukey mutiple comparison *post-hoc* testing. MIA-treated rats had lowered ipsilateral PWT in both strains, though this occurred earlier and to a greater extent in WKY rats when compared to Wistar rats (**D**). Data are mean ± SEM change in vFH compared to baseline, * p<0.05, ** p<0.01, versus WKY saline, # p<0.05, ## p<0.01 versus Wistar saline, + p<0.05, versus Wistar MIA, 2 way ANOVA with Tukey multiple comparison *post-hoc* testing. Measurement of anxiety-like behaviour in the EPM at D21 after model induction revealed no significant differences between MIA- and saline-injected rats for the two strains (**E**), but there remained a significant anxiety-like phenotype in the WKY strain when compared to the Wistar strain (**F**). Data represent AUC of time spent in the open arm for each minute of the 10 min trial, individual data points are shown with bars indicating the median value for each group, and error bars representing the IQR. ## p<0.01 versus Wistar saline, Kruskal-Wallis test with Dunn’s multiple comparison *post-hoc* test, ++++ p<0.001 versus Wistar rats, one-tailed Mann-Whitney *U* test. To ensure any behavioural differences observed in the EPM did not result from strain differences in locomotion, locomotor activity was assessed over a 1 hour period at baseline, and 18-21 days after model induction (**G**). No significant differences between strains were observed at either time point. Data are expressed as mean ± SEM beam breaks per hour, adjusted for bodyweight. Macroscopic assessment of cartilage damage in ipsilateral knee joints revealed extensive widespread cartilage damage in MIA-injected rats of both strains, with little or no damage observed in saline-injected animals (**H**). Data represent the summed scores for each of the 5 individual joint compartments (0-5, max score 25), individual data points are shown, with bars representing the median value and error bars the IQR. #### p<0.001 versus Wistar saline, **** p<0.001 versus WKY saline.

#### 2.7.2 Spinal electrophysiology

Degree of wind up was determined from the total number of spikes recorded in response to each stimuli in a train of 16 delivered at 3 x C-fibre threshold. Responses corresponding to each fibre type were binned according to post-stimulus latency, and wind up calculated from those falling in the C-fibre (90-300ms) and post-discharge (300-1000ms) latency range. Responses to mechanical stimuli were recorded as average firing rate (Hz) in response to 10s stimulation with each vFH. To determine effects of pharmacological interventions, mean maximal inhibition (MMI) for each dose was calculated as maximal % change versus baseline, with peak inhibition occurring 40-60 mins after administration. MMI data for each dose were then plotted for each strain and treatment, and the AUC calculated and compared.

#### 2.7.3 Statistical Analyses

Group sizes were calculated based on previous similar studies using the MIA model to give sufficient statistical power whilst minimising animal usage. Data were analysed using Prism 8.3.1 software (GraphPad, La Jolla, USA). Data distributions were assessed via D’Agostino & Pearson normality testing, and subsequently treated as parametric or non-parametric, as appropriate. Differences in pain behaviour data were analysed via two-way ANOVA, with strain and treatment as the independent variables and Tukey’s *post-hoc* test for multiple comparisons. For all other comparisons one-way ANOVAs with Holm-Sidak multiple comparisons, or Kruskal-Wallis tests with Dunn’s multiple comparison *post-hoc* tests were used to analyse data with 3 or more groups, whilst Mann-Whitney *U* tests or Wilcoxon signed rank tests were used for comparisons between strains. Data are stated as mean ± SEM, or median with interquartile range (IQR), as appropriate.

Full experimental data are available from the authors upon request.

### 3. Results

#### 3.1 WKY rats exhibit a basal anxiety-like phenotype

To confirm the anxiety-like phenotype of WKY rats, behaviour in the anxiogenic environment of the elevated plus maze (EPM) was assessed in both strains at baseline. WKY rats spent significantly less time in the open arms when compared to Wistar rats (Figure 1A, median AUC 17.40, IQR 4.92 – 31.68 versus 33.60, IQR 19.2 – 57.87, p=0.003), and had a significantly greater average latency to enter the open outer arm (600 versus 298, IQR 86-600 seconds, p=0.0077, Mann-Whitney *U* test, Figure 1B). Collectively, these data confirm that the WKY rats used in this study had an anxiety-like behavioural phenotype.

#### 3.2 WKY rats exhibit exacerbated pain behaviour in the MIA model of OA

WKY and Wistar rats had comparable hindpaw mechanical sensitivity at baseline and weight was borne equally on both hindpaws (Table 1). Consistent with our previous work, intra-articular injection of MIA resulted in a significant shift in weightbearing from the injured side (weightbearing asymmetry) in both strains of rats from day 3 onwards, which persisted for the duration of the study (Figure 1C). The degree of asymmetry following injection of MIA was similar in both strains of rats, although slightly less pronounced in WKY rats. Both strains of rats also exhibited a lowering of the ipsilateral mechanical hindpaw withdrawal threshold (PWT, Figure 1D) which was evident from day 10. Ipsilateral PWTs were lowered to a greater extent in the WKY strain when compared to Wistar rats (Table 1; mean change in vFH from baseline Wistar = −1.68, WKY = −3.22), indicating an exacerbated OA-like pain phenotype, as previously reported (Burston et al., 2019). We also observed a reduction in contralateral PWTs in WKY rats (**Extended Data Figure 1-1**), which is consistent with a centralised pain phenotype in this model (Burston et al., 2019).

**Table 1.**
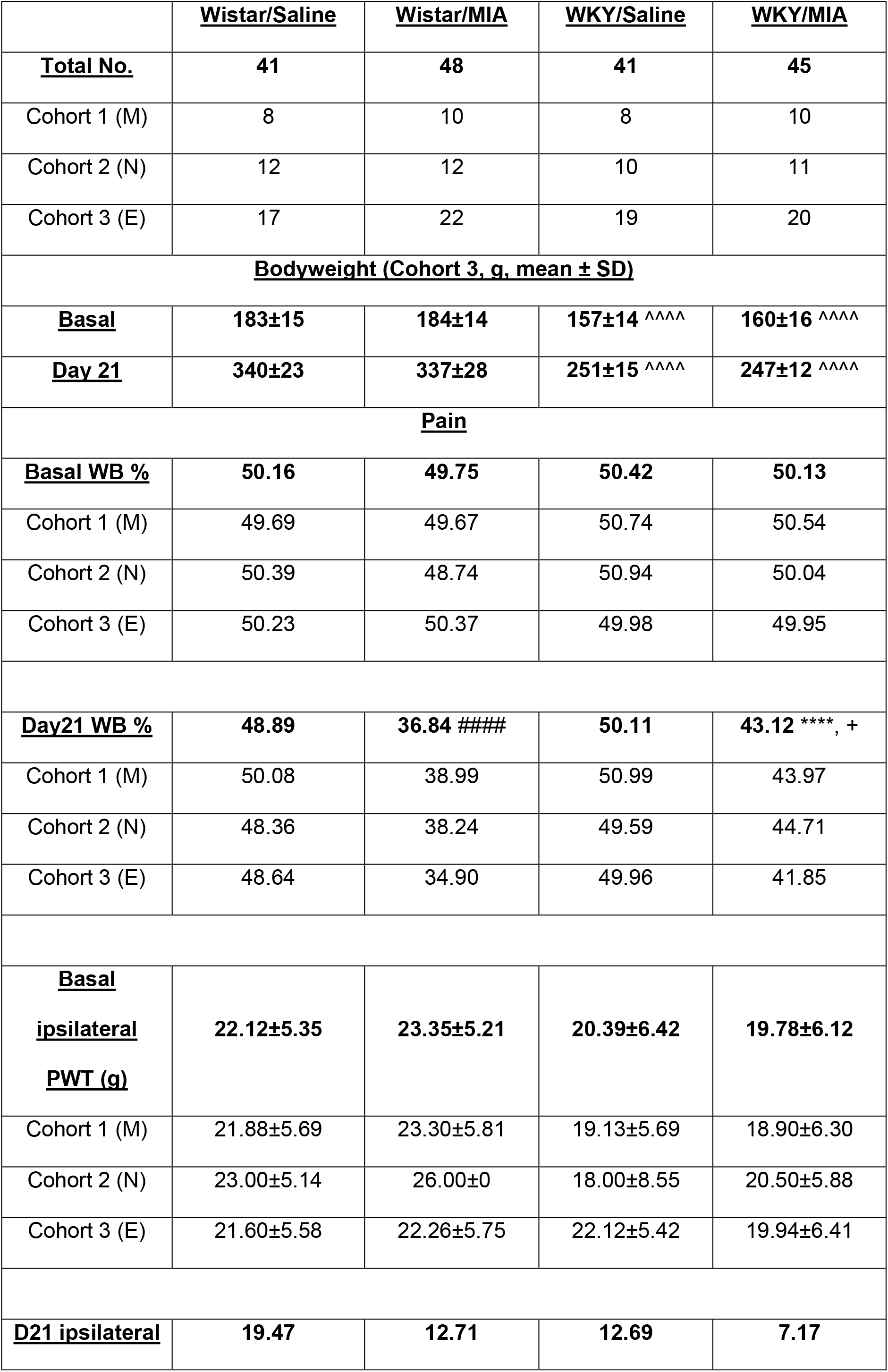
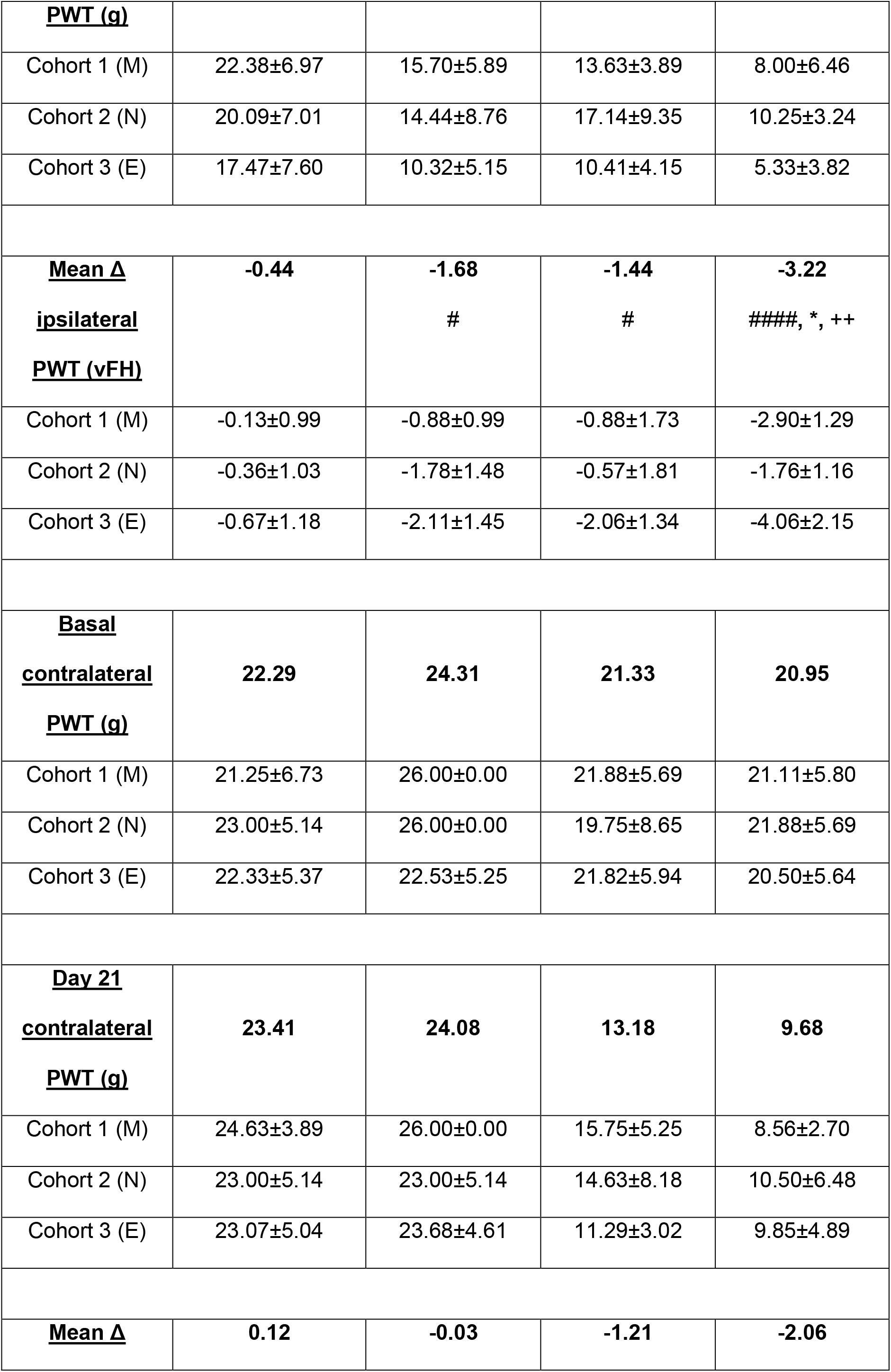
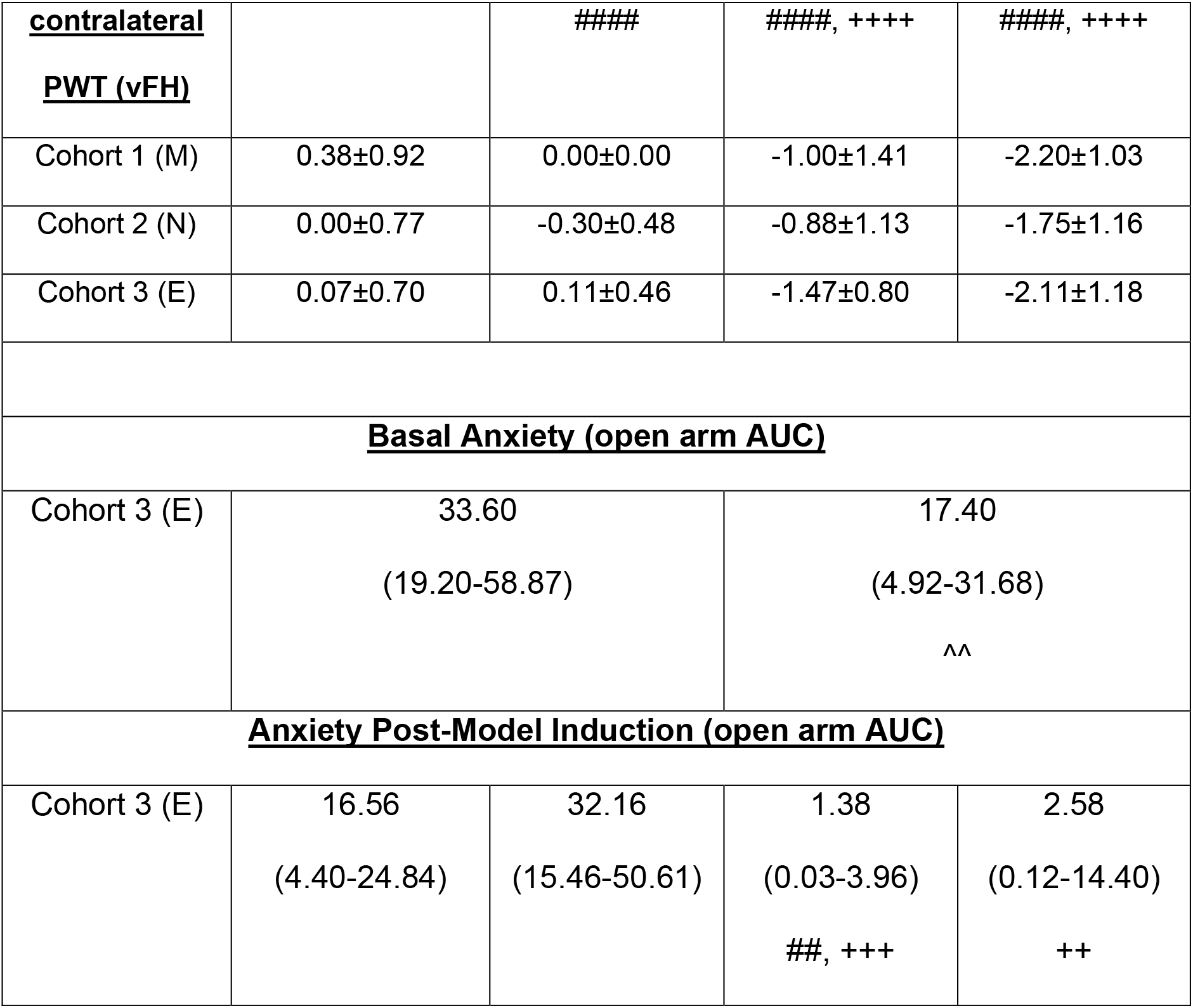
Summary of animal numbers and behavioural data. Data from all 3 cohorts of animals utilised in this study (M = morphine study, N = naloxone study, E = electrophysiology study). Data are expressed as mean ± SD, or median (IQR). Significance assessed via 2 way ANOVA with Tukey multiple comparison *post-hoc* testing, Kruskal-Wallis test with Dunn’s multiple comparison *post-hoc* testing, or Mann-Whitney *U* tests as appropriate. ^^ p<0.01, *^^^^* p<0.0001 versus Wistar/saline & Wistar/MIA # p<0.05, #### p<0.0001 versus Wistar/saline * p<0.05, **** p<0.0001 versus WKY/saline + p<0.05, ++ p<0.01, +++ p<0.001, ++++ p<0.0001 versus Wistar/MIA.

#### 3.3 Anxiety-like behaviour and OA-like joint pathology

To determine whether induction of OA-like pain behaviour altered anxiety-like behaviours, behavioural responses in the EPM were reassessed 19-21 days after intra-articular injection of saline or MIA. No significant differences were observed between MIA and saline-injected rats when comparing within strains (Figure 1E), however an anxiety-like phenotype was present in the WKY strain when compared to Wistar rats at this time point (Figure 1F, AUC for time spent in open arms 1.50 0.03 – 6.48 versus 20.10 IQR 5.98 – 41.61, **** p<0.0001, one-tailed Mann Whitney U test). This is in agreement with our previous work demonstrating that presence of OA-like pain behaviour does not further increase anxiety-like behaviour in either strain (Burston et al., 2019). Locomotor activity at baseline and at D18-21 was comparable between the two strains of rats (Figure 1G), demonstrating no differences in exploratory activity in the absence of an anxiogenic environment.

To ensure the marked increase in MIA-induced pain behaviour in the WKY strain was not due to an alteration in joint pathology, cartilage damage in the injected knee was assessed via macroscopic scoring. MIA administration was associated with significant increases in cartilage damage compared to the appropriate saline control, in both WKY and Wistar rats (Figure 1H). It is noteworthy that, despite the significantly greater pain behaviour in MIA-treated WKY rats, this strain actually had slightly less cartilage damage when compared to Wistar rats.

#### 3.4 Exacerbated pain behaviour in the WKY/MIA model is associated with enhanced excitability of spinal neurones

To determine whether the altered behavioural phenotype in the MIA model of OA-like pain behaviour in WKY rats was encoded in the spinal cord dorsal horn, a key site in neuronal nociceptive circuitry, we then performed *in vivo* single-unit recordings of spinal WDR neurons in both WKY and Wistar rats 21-27 days after model induction. No differences were observed in baseline characteristics of recorded neurones (depth, Aβ and C-fibre latencies and thresholds, see **Extended Data Table 2-1**). However, wind up, a proxy of central sensitization (Li et al., 1999), did significantly vary across strains and treatments (Figure 2A). In the absence of a pain state, wind-up was significantly greater in WKY rats when compared to their Wistar counterparts (p<0.05 for stimuli 5 & 6, 2 way ANOVA with Tukey multiple comparisons *post-hoc* testing), and was further increased 21 days after MIA-treatment in the WKY strain (p<0.05, stimuli 3-6, 2 way ANOVA with Tukey multiple comparisons *post-hoc* testing). The response to the first stimulus in the train was also notably larger in the WKY strain (Figure 2A), particularly responses occurring at Aδ and C fibre latencies in WKY rats (**Extended Data Figure 2-2**). Despite a marked lowering of ipsilateral PWTs in the MIA model in both strains, there were no differences in the magnitudes of innocuous and noxious mechanically-evoked responses of WDR neurons between any of the groups (Figure 2B). Taken together, these data demonstrate enhanced spinal sensitivity to nociceptive input in the WKY strain, with further enhanced excitability to overt nociceptive stimuli in the presence of an OA-like pain state.

**Figure 2.**
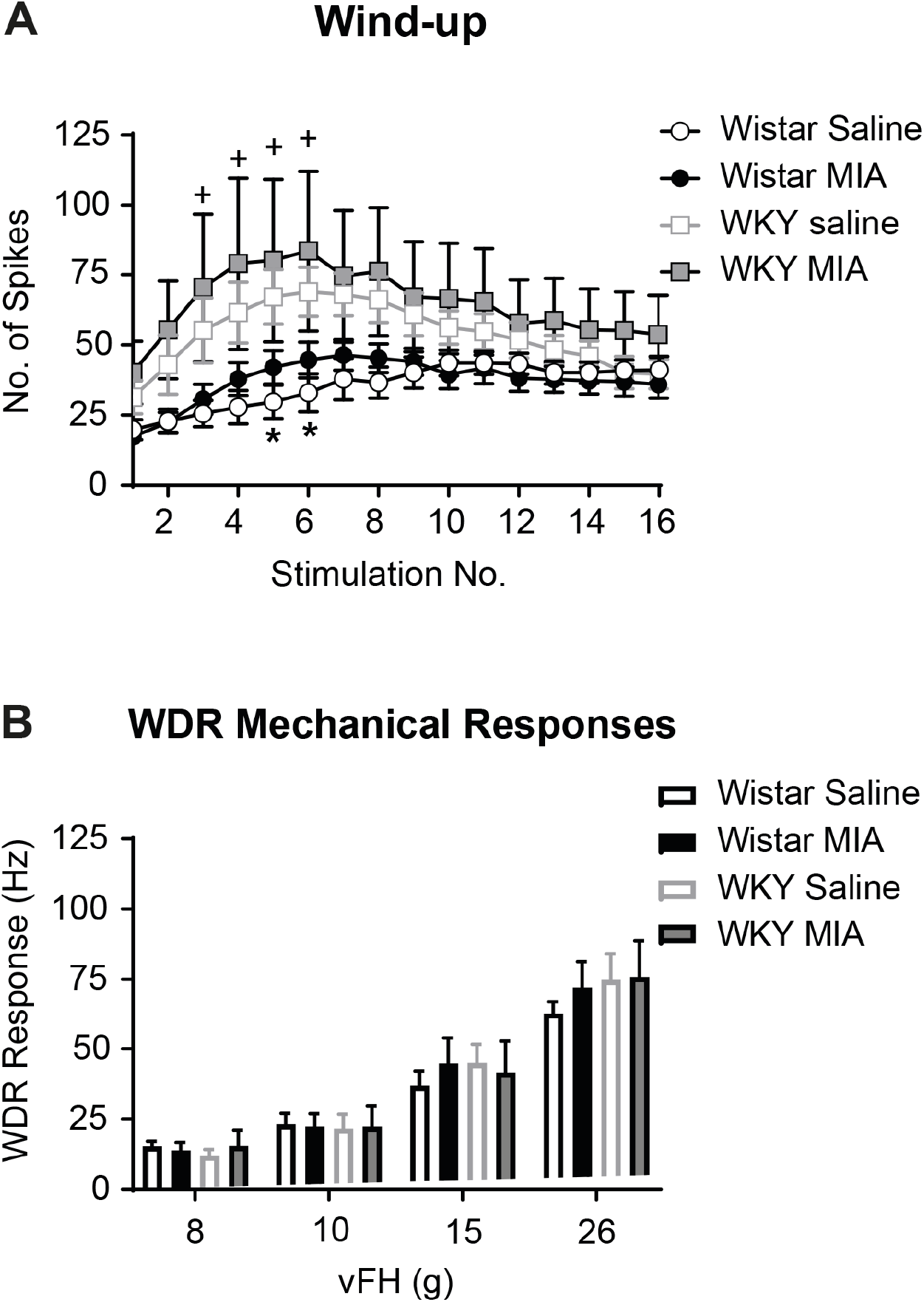
Enhanced spinal excitability and nociceptive responses in the WKY-MIA model. *In vivo* electrophysiological recordings of WDR neurons in the deep DH revealed enhanced wind up in the WKY strain in response to noxious electrical stimulation at 3 x C-fibre threshold (**A**). The number of action potentials recorded in the 90-1000ms post-stimulus (C-fibre to post-discharge latency range) was significantly higher in saline-injected WKY rats (stimuli 5 & 6) and MIA-injected WKY rats (stimuli 3-6) when compared to their Wistar counterparts. Data represent mean ± SEM number of action potentials. * p<0.05 versus WKY saline, + p<0.05 versus Wistar MIA, 2 way ANOVA with Tukey multiple comparisons *post-hoc* testing. There were no significant differences in the frequency of WDR firing recorded in response to mechanical stimulation across a range of vFHs between strains or treatments (**B**). Data represent the frequency of firing in response to a 10s stimulus with each vFH. Values are means ± SEM.

#### 3.4 Reduced effects of systemic morphine in the WKY/MIA model of high anxiety & OA-like pain behaviour

At 21 days following induction of the MIA model of OA pain, subcutaneous (s.c.) injection of the opioid receptor agonist morphine (0.5, 2.5, & 6 mg.kg^-1^ cumulative dose) produced a significant, dose-related reduction in weightbearing asymmetry in Wistar rats (Figure 3A). In contrast, only the highest dose of morphine significantly inhibited weightbearing asymmetry in MIA-treated WKY rats (Figure 3A). Similarly, morphine had a significantly blunted effect on ipsilateral PWTs in MIA-treated WKY rats when compared to the Wistar strain (Figure 3B). All three doses of morphine significantly reversed PWTs in MIA-treated Wistar rats, whereas only the highest dose of morphine produced a significant reversal in ipsilateral PWTs in WKY rats (Figure 3B). This highest dose of morphine significantly reversed the lowered contralateral PWT evident in MIA treated WKY rats (Figure 3C).

**Figure 3.**
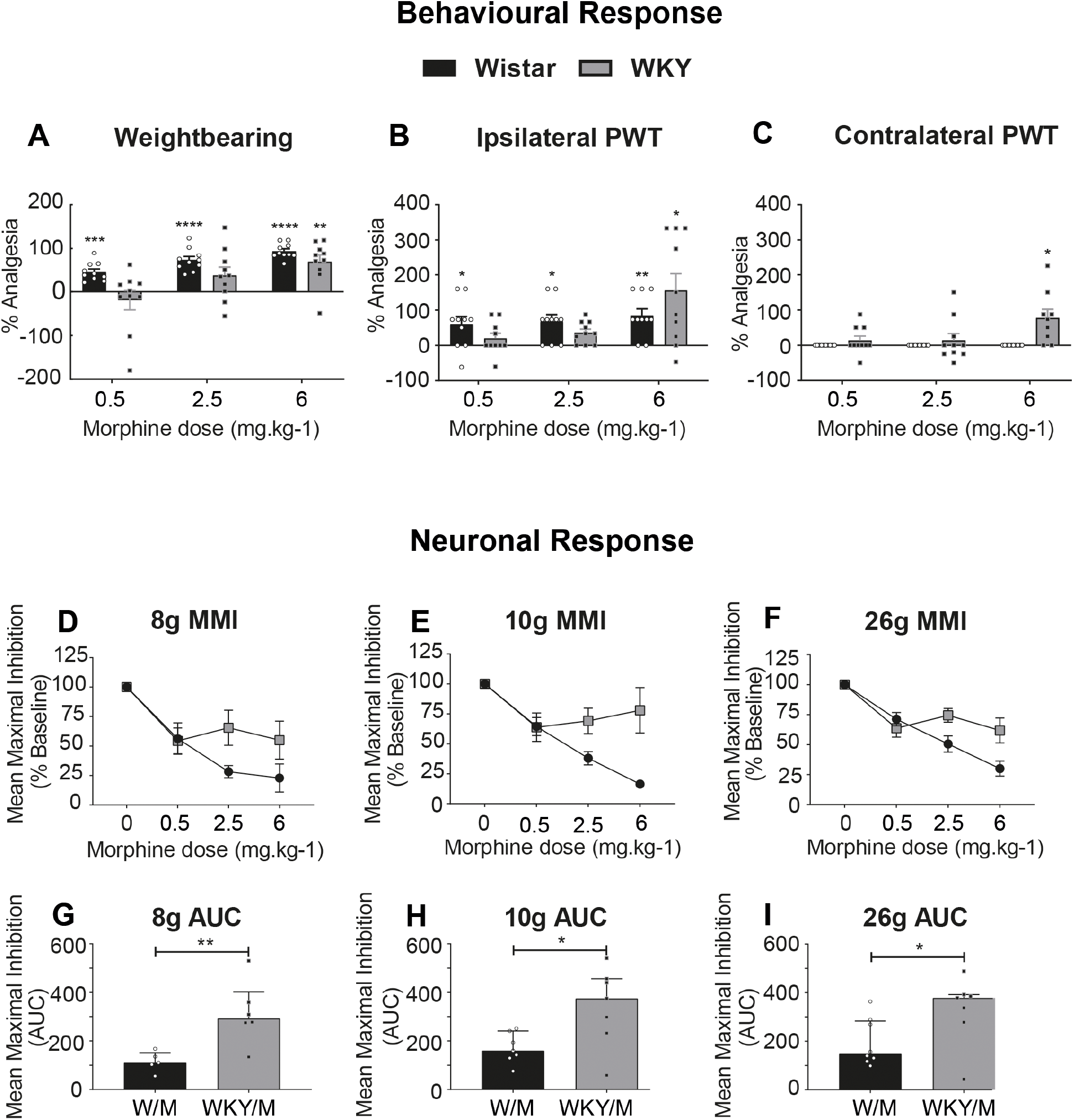
Reduced behavioural & spinal response to systemic morphine in the WKY/MIA model of high anxiety & OA-like pain. Systemic administration of morphine produced dose-related inhibitions in the MIA model in Wistar rats. By contrast, inhibitory effects of the low dose of morphine were significantly attenuated in the WKY strain for weightbearing asymmetry (**A**) and ipsilateral PWTs (**B**). Morphine did not significantly alter contralateral PWTs in MIA-treated Wistar rats. However, 6 mg.kg^-1^ morphine reversed lowered contralateral PWTs in MIA-treated WKY rats (**C**). Data represent % analgesia to 3 cumulative doses of systemic morphine, with abolition of weightbearing asymmetry representing 100% analgesia in **A**, and a return to PWTs to pre-model basal values representing 100% analgesia in **B** & **C**. * p<0.05, ** p<0.01, ***p<0.001, ****p<0.0001, Wilcoxon Signed Ranks test with a hypothetical value of 100. Effects of systemic morphine on mechanically-evoked responses of spinal WDR neurones was also blunted in MIA treated WKY rats. There was a blunted effect of cumulative doses of 2.5 and and 6 mg.kg^-1^ morphine on 8g- (**D**), 10g- (**E**), and 26g-evoked (**F**) vFH evoked responses of WDR neurones in MIA-treated WKY rats, which was confirmed by AUC analysis (**G**-**I**). Data represent median ± IQR, * p<0.05, ** p<0.01 Mann-Whitney *U* tests.

At the level of the spinal cord, the inhibitory effect of cumulative systemic (s.c.) doses of morphine on evoked neuronal responses was also significantly blunted in MIA-treated WKY rats when compared to Wistar rats (Figure 3D-I). Although the mean maximal inhibitory (MMI) effect of the lowest dose (0.5 mg.kg^-1^) was similar across the range of noxious and non-noxious mechanical stimuli studied, there was little additional inhibition following successive increased doses of morphine in the WKY MIA rats. This was particularly evident with the lower force filaments (8 & 10g, Figure 3D-E, & G-H). These data provide further evidence of reduced inhibitory effects of systemic morphine in rats with an anxiety-like phenotype and OA-like pain.

#### 3.5 Evidence for altered endogenous opioid signalling in WKY rats

We hypothesised that the reduced inhibitory effects of morphine on pain behaviour and spinal neuronal responses in WKY rats may arise as a result of changes in opioid receptor function and circulating levels of endogenous opioids. To test this, plasma levels of β-endorphin were measured by ELISA. Significantly higher levels of β-endorphin were detected in WKY rats when compared to Wistar rats (Figure 4A). We then focused on MOR expression at the level of the dorsal horn of the spinal cord. Western blotting revealed no significant differences in total spinal MOR expression in any group (Figure 4B&C). There was, however, an increase in the proportion of MOR phosphorylated at the master phosphorylation site (serine residue 375; P-ser375) in the dorsal horn of the spinal cord in MIA-treated WKY rats, when compared to saline-treated WKY rats (Figure 4D, p = 0.0143). By contrast, the proportion of phosphorylation at P-ser375 in the dorsal horn of the spinal cord was not altered in MIA-treated Wistar rats compared to saline-treated Wistar rats, and levels were comparable in the saline-treated WKY rats.

**Figure 4.**
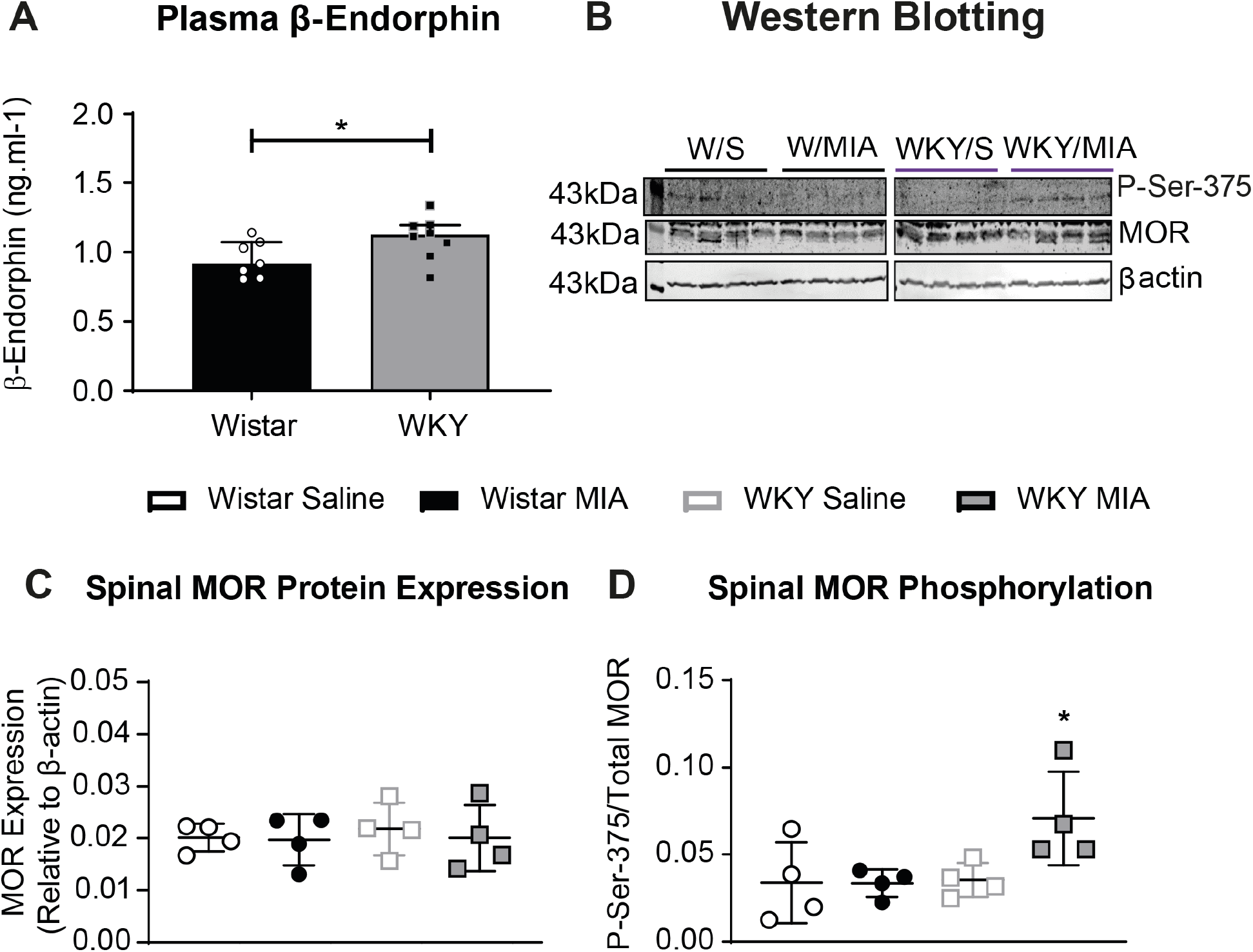
Altered endogenous opioid function in the WKY/MIA model. Assessment of circulating plasma levels of β-endorphin in terminal blood samples from naïve Wistar (*n* = 7) and WKY (*n* = 8) rats revealed a 20% increase in the WKY strain (**A**). Bars represent the median value, error bars indicate IQR. * p = 0.0361, one-tailed Mann-Whitney *U* test. Western blotting depicting expresssion of total MOR, and MOR phosphorylated at the master phosphorylation site (P-ser375) in whole lumbar spinal cord homogenates (**B**). Densitometry quantification revealed no change in total MOR levels (**C**) across strains and treatments (*n* = 4/group), and no significant change in phosphorylation in MIA-treated Wistar rats (**D**). However, a significant increase in the proportion of P-ser375-MOR was evident in MIA-treated WKY rats when compared to saline-treated controls. Lines at median values, error bars represent IQR, * p = 0.0143, one-tailed Mann-Whitnety *U* test.

As our data so far point towards altered opioidergic function in WKY rats, the effects of blocking the μ-opioid receptor with the antagonist naloxone on pain behaviour were assessed in the MIA model in both strains of rats (Figure 5). Naloxone (0.1-1 mg.kg^-1^ s.c.) did not alter PWTs in Wistar rats 21 days after intra-articular injection of saline, suggesting no overt basal endogenous opioidergic tone in these rats. However, all three doses of naloxone (0.1-1 mg.kg^-1^) significantly lowered ipsilateral PWTs in MIA-treated Wistar rats, supporting the presence of endogenous opioid tone following induction of the model of OA pain in this strain of rats. In WKY rats, systemic naloxone (0.1-1 mg.kg^-1^) produced a significant bilateral lowering of PWTs (Figure 5A-D) which was similar in both saline- and MIA-treated WKY rats. These data further support the presence of an elevated endogenous opioid tone in WKY rats in the absence of the pain model, consistent with the increased levels of β-endorphin demonstrated herein. The endogenous opioid tone did not appear to be further increased in the presence of the model of OA pain.

**Figure 5.**
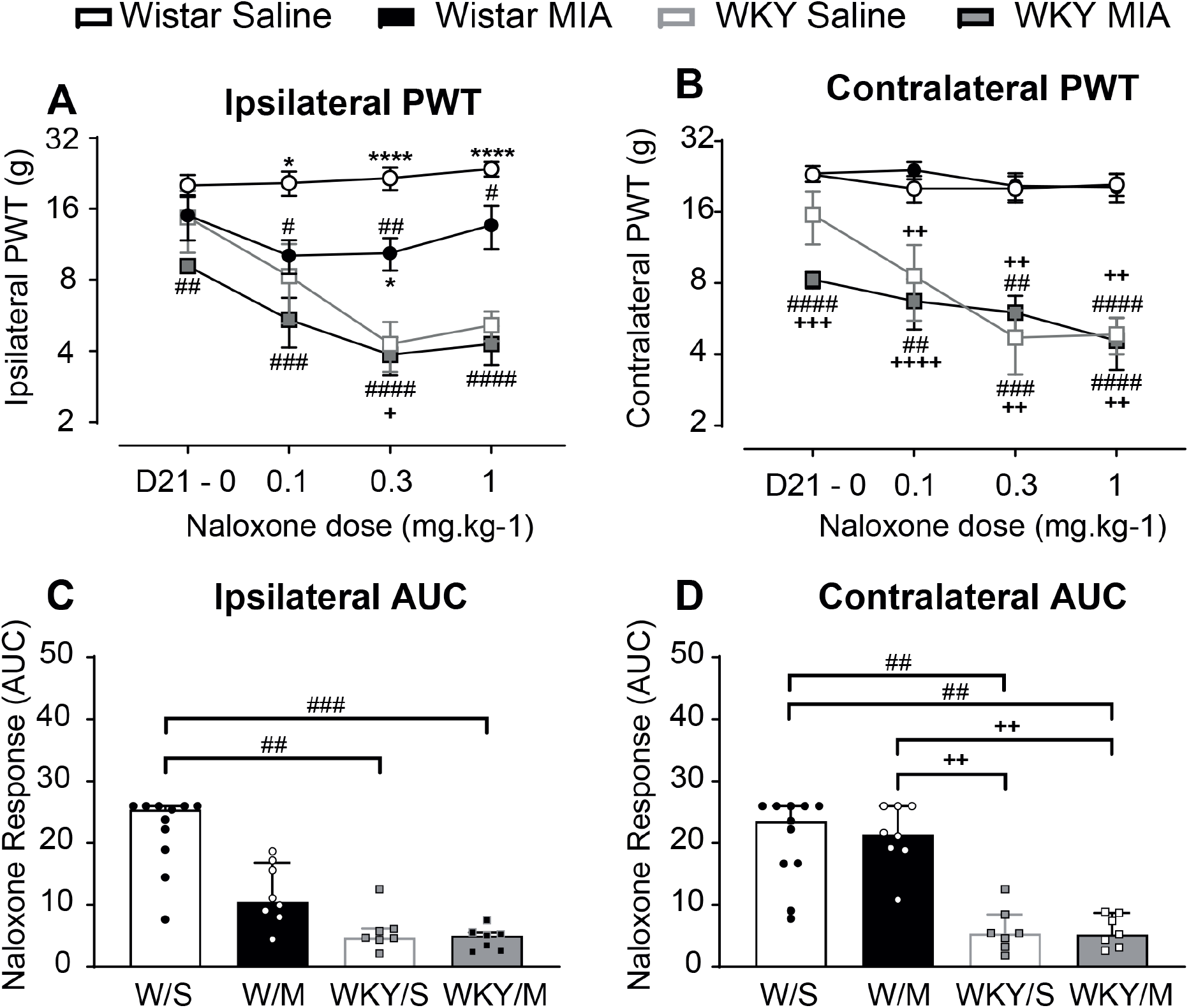
Altered endogenous opioid tone in rats with an anxiety-like phenotype. Systemic administration of the MOR antagonist naloxone (0.1-1 mg.kg^-1^, s.c.) significantly reduced ipsilateral PWTs in MIA-treated, but not saline-treated, Wistar rats (**A**), suggesting elevated opioidergic tone in the model of OA-like pain. In WKY rats, naloxone produced a significant, dose-dependent, and bilateral reduction in PWTs in both saline- and MIA-treated WKY rats (**A&B**), suggesting a more generalised elevation of opioidergic tone in this strain. Data represent mean ± SEM for baseline PWTs, and MMI values for each cumulative dose of naloxone. # p<0.05, ## p<0.01, ### p<0.001, #### p<0.0001 versus Wistar saline; *p<0.05, **** p<0.0001 versus WKY saline; + p<0.05, ++p<0.01, +++ p<0.001, ++++ p<0.0001 versus Wistar MIA, 2 way ANOVA with Tukey multiple comparison *post-hoc* testing. AUC analysis of dose response curves revealed a significantly greater effect of naloxone on ipsilateral (**C**) and contralateral (**D**) PWTs in WKY rats when compared to Wistar rats. Individual data points are shown with bars representing median values and error bars depicting IQR. ## p<0.01, ### p<0.001 versus Wistar saline; ++p<0.01 versus Wistar/MIA, Kruskal-Wallis test with Dunn’s multiple comparison *post-hoc* testing.

### Discussion

Clinically, people with high anxiety and chronic pain, including OA pain, have greater opioid consumption compared to people with comparable pain but less anxiety (Barnett et al., 2018). Here, we demonstrate reduced morphine-mediated analgesia in a rodent model of heightened anxiety and OA-like pain, providing an experimental model to further investigate the underlying neurobiological mechanisms.

Systemically-administered morphine produced a robust inhibition of behavioural pain responses following OA-induced joint pathology in Wistar rats. However, in WKY rats with a heightened anxiety-like phenotype, the effects of morphine on OA-induced pain behaviour were significantly blunted. In particular, the lowest dose of morphine significantly reversed weightbearing asymmetry and hindpaw withdrawal thresholds in OA-treated Wistar, but not WKY, rats. Since the highest dose of systemic morphine produced comparable inhibitory effects on MIA-induced pain behaviour in both strains of rats, it is likely that this results from reduced efficacy of opioid signalling in WKY rats. Reduced efficacy of morphine was previously reported in acute pain tests in WKY rats (Hestehave et al., 2019). We provide the first evidence of blunted opioid analgesia in a clinically-relevant model of chronic OA pain in this genetic strain of rats, providing a platform to interrogate the underlying neurobiology. Our experimental data are consistent with clinical evidence that anxiety in people with joint pain is associated with the greater use of prescription opioids (Barnett et al., 2018), and prolonged use of opioids following total joint replacement for persistent joint pain (Namba et al., 2018). Catastrophizing has also been associated with increased prescription opioid use 4 years following total joint replacement (Valdes et al., 2015). Interestingly, recent work reported that the relationship between anxiety and prescription opioid use for chronic pain is stronger in males than females (Rogers et al., 2020), supporting our strategy of using male rats for this preclinical experimental model system.

Electrophysiological recordings of spinal cord dorsal horn WDR neurones in OA-treated WKY rats compared to OA-treated Wistar rats also revealed subtle changes in MOR function. Specifically, the effects of systemic morphine on WDR responses to low- and high-force mechanical stimulation of the hindpaw were blunted in WKY OA-treated rats, supporting the notion that MOR function is altered in these animals. Clearly, these changes in morphine responsiveness may reflect alterations in MOR function at multiple levels of the neuraxis. We hypothesised that these blunted inhibitory effects of exogenous morphine may arise due to a loss or desensitization of MOR.

The C-terminus of MOR has a number of phosphorylation sites that contribute to receptor desensitization and internalisation. Of the residues that undergo agonist-dependent phosphorylation, residues 375 to 379 (STANT) have a critical role in endogenous opioid-induced acute desensitization, recovery from desensitisation, and internalisation of MOR (Arttamangkul et al., 2019; Kliewer et al., 2019). In our study, phosphorylation of the serine-375 residue of MOR was significantly elevated in the ipsilateral dorsal horn of the spinal cord of OA-treated WKY rats. These data suggest that MOR tolerance in the dorsal horn spinal cord may account, at least in part, for the loss of analgesic effect of morphine in these rats. However, the spinal cord is unlikely to be the only site and changes in MOR expression and/or phosphorylation at other sites in the brain may also contribute.

Previous studies have reported complex relationships between endogenous opioid function, depressive symptoms, and trait anxiety in people (Burns et al., 2017). In our study, a small, but significant, elevation in plasma β-endorphin was detected in WKY rats. This finding, alongside the demonstration of a reduced analgesic effect of systemic morphine in the model of OA pain, is consistent with the clinical evidence that greater endogenous opioid function is associated with lower morphine analgesic responsiveness (Bruehl et al., 2013). A limitation of our study is that plasma levels of β-endorphin were not measured in MIA-treated WKY rats with OA-like pain, nor were levels of other endogenous opioids determined. Nevertheless, based on previous studies of the MIA model in mice (Aman et al., 2019), we predict that β-endorphin levels are likely further elevated in the model of OA pain.

Functional evidence for changes in endogenous opioid tone in WKY rats was provided by our demonstration that systemic naloxone lowered hindpaw withdrawal thresholds in Wistar rats in the presence of the model of OA pain, but not in pain free saline-treated Wistar rats. These data are consistent with the engagement of the endogenous opioidergic systems in models of chronic pain previously demonstrated in clinical pain states (Levine et al., 1978; Kayser and Guilbaud, 1990; reviewed in Fields, 2004). Under these conditions, the endogenous opioidergic systems acts to counter the increased pro-nociceptive signalling and the manifestation of pain behaviour and experience. In the WKY strain, naloxone significantly lowered hindpaw withdrawal thresholds in the absence of the model of OA pain, consistent with the elevated levels of plasma β-endorphin in WKY rats demonstrated herein. The presence of the model of OA pain in WKY rats did not further increase the effect of naloxone on hindpaw withdrawal thresholds, suggesting that the presence of the model of OA pain could not further engage the endogenous opioidergic system in the WKY rats, as it did in the Wistar rats. This loss of chronic pain-induced engagement of the opioidergic inhibitory pathways, alongside our evidence for increased phosphorylation of serine-375 of MOR in WKY MIA treated rats, may account for the reduced effectiveness of the lower doses of morphine in WKY MIA treated rats.

A consideration of this work is the use of an inbred rat strain to model anxiety-like behaviour, and whether this may confound our findings. Differences in MOR gene expression between WKY and Sprague Dawley rats have been reported for some brain regions (Burke et al., 2019). Although no differences in expression were observed in the reward-associated nucleus accumbens (Dennis et al., 2016), and the report of reduced acquisition of morphine-induced conditioned place preference in WKY rats, supports our evidence for dysfunctional responses to exogenous opioids in this strain. Alongside altered opioid function, WKY rats have lower basal levels of limbic serotonin and dopamine, resulting in a blunted serotonergic and noradrenergic response to acute stress (De La Garza and Mahoney, 2004; Yamada et al., 2013) which may be relevant to the clinical situation. Given the key roles for supraspinal monoamines in descending modulation of pain signalling (Bannister and Dickenson, 2016), it is likely that both opioidergic and monaminergic dysfunction contribute to augmented OA-like pain responses in WKY rats.

Our study highlights the functional significance of the combination of anxiety, chronic pain, and elevated opioidergic tone, which leads not only to increased pain behaviour, but also decreased efficacy of opioid analgesia. Broader implications of our work include caution in the prescription of opioids to manage chronic pain in OA patients with comorbid high anxiety.

## Acknowledgements

This work was supported by Arthritis Research United Kingdom (grants 18769, 20777) and the Medical Research Council (studentship 1653552). The authors wish to thank Rob Lane (University of Nottingham) for helpful discussions and advice on opioid receptor pharmacology.

