## Extended data for "Anxiety enhances pain in a model of osteoarthritis and is associated with altered endogenous opioid function and reduced opioid analgesia"

**Extended Data Table 1-1 – Animals excluded from study**

| **1. Non-matching cartilage scores** | **Wistar/Saline** | **Wistar/MIA** | **WKY/Saline** | **WKY/MIA** |
|---|---|---|---|---|
| **Total No.** | **3** | **4** | **3** | **4** |
| Cohort 1 (M) | 0/8 | 0/10 | 0/8 | 0/10 |
| Cohort 2 (N) | 1/12 | 1/12 | 1/10 | 2/11 |
| Cohort 3 (E) | 2/17 | 2/22 | 3/19 | 2/20 |
| **2. Incomplete behavioural data** | **Wistar/Saline** | **Wistar/MIA** | **WKY/Saline** | **WKY/MIA** |
| Cohort 2 (N) | 1/12 | 1/12 | 1/10 | 2/11 |
| **3. Incomplete electrophysiology data** | **Wistar/Saline** | **Wistar/MIA** | **WKY/Saline** | **WKY/MIA** |
| Cohort 3 (E) | 2/10 | 5/15 | 3/12 | 2/12 |

**M = morphine study, N = naloxone study, E = electrophysiology study**

**1.** A total of 14 rats were excluded from the study on the basis of joint pathology inconsistent with the recorded intra-articular treatment received. The exclusion criteria were any rats with a total cartilage damage score < 6 for MIA-treated groups, or ≥ 6 for saline-treated groups. These discrepancies likely resulted from experimenter error during blinding, or misplacement of intra-articular injection.

**2.** Behavioural data from 4 rats were excluded from the naloxone time course study due to environmental noise disruption, preventing the collection of valid pain behaviour.

**3.** Incomplete electrophysiological datasets were obtained from 12 animals due to loss of the target cell during recordings

**Extended Data Figure 1-1 - Contralateral pain phenotype in the WKY-MIA model**

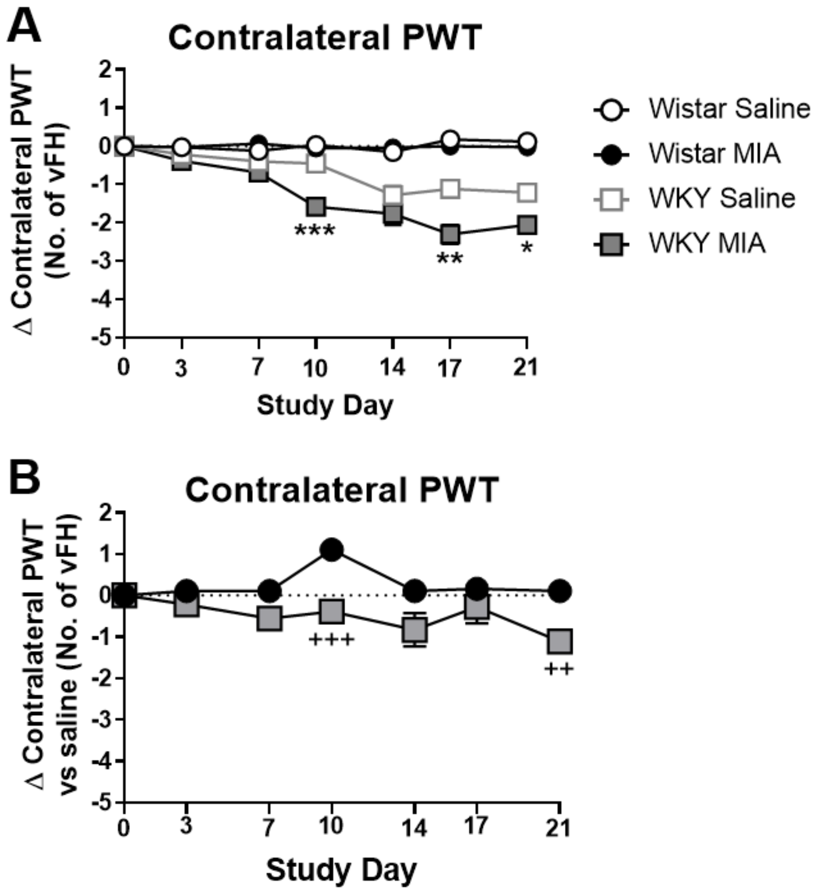


Injection of MIA or saline did not alter PWTs in the contralateral hindlimb in Wistar rats, but did lower contralateral PWTs in the WKY strain (**A**). Normalising data from each strain to their respective saline-treated controls revealed a significantly greater decrease in contralateral PWTs in the WKY strain after MIA injection (**B**).

Data are mean ± SEM change in vFH compared to baseline, * $p<0.05$, ** $p<0.01$ *** $p<0.001$ versus WKY saline, + $p<0.05$, +++ $p<0.001$ versus Wistar MIA, 2 way ANOVA with Tukey multiple comparison *post-hoc* testing.

**Extended Table 2-1 – WDR neuron characteristics**

| | Depth (μm) | Aβ Fibre Threshold (mA) | Aβ Fibre Latency (ms) | C Fibre Threshold (mA) | C-Fibre Latency (ms) |
|---|---|---|---|---|---|
| **Wistar Saline** | 770 (630 – 873) | 0.13 (0.11 – 0.15) | 9 (6 - 12) | 1.00 (0.80 – 1.38) | 184 (144 - 243) |
| **Wistar MIA** | 775 (650 – 893) | 0.14 (0.11 – 0.15) | 11 (7 - 12) | 1.00 (0.90 – 1.10) | 205 (167 - 241) |
| **WKY Saline** | 845 (630 - 980) | 0.10 (0.09 – 0.14) | 6 (4 - 12) | 1.00 (0.90 – 1.10) | 144 (108 - 195) |
| **WKY MIA** | 780 (745 - 805) | 0.11 (0.09 – 0.13) | 6 (6 – 11) | 1.00 (0.88 – 1.13) | 223 (157 – 271) |

No significant differences were observed in the depth, or Aβ or C-Fibre thresholds or latencies between experimental groups in this study. There was, however, a slight trend towards decreased Aβ latencies and thresholds in the WKY strain, and increased C-fibre latencies in cells from MIA-treated rats of either strain when compared to saline-treated controls.

Data are median values with IQR. Statistical comparisons via Kruskal-Wallis test with Dunn’s multiple comparison *post-hoc* testing.

**Extended Data Figure 2-2 - Enhanced spinal responses to nociceptive input in the WKY-MIA model**

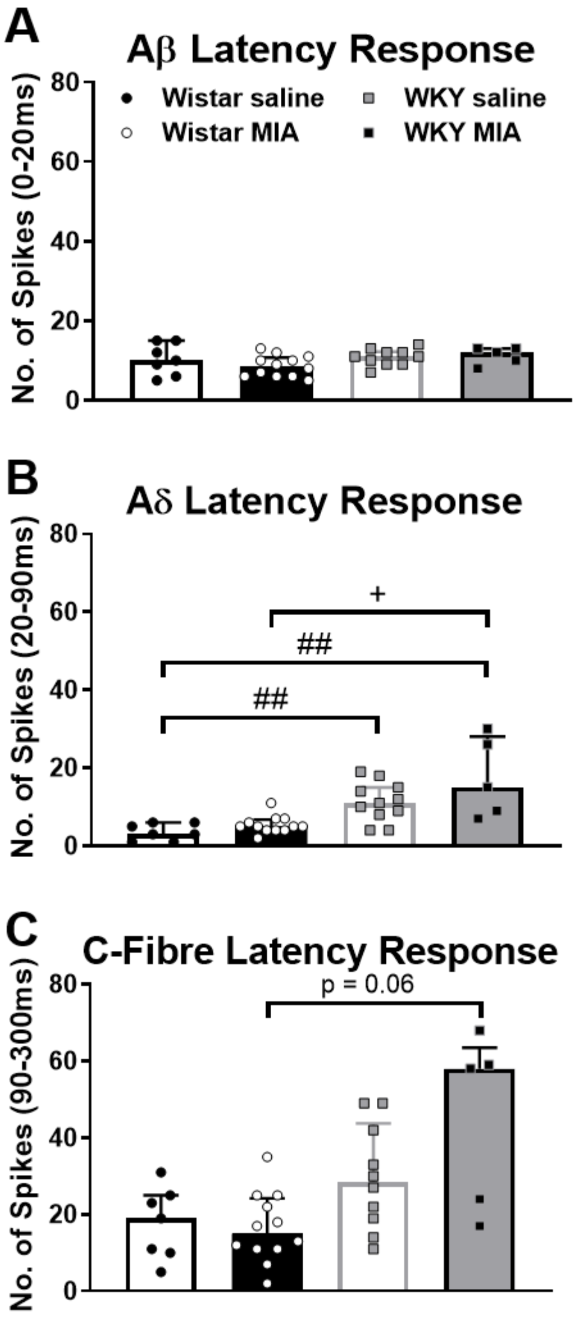


WDR responses to electrical stimulation at 3 x C fibre threshold binned by response latency. Responses in the Aβ fibre latency (0-20ms) were similar across strains and treatments (**A**), whilst action potentials in the Aδ fibre latency range were 3-fold higher in the WKY strain when compared to the Wistar strain (**B** Wistar saline = 3, IQR 4-7 versus WKY saline = 11, IQR 8-15, p=0.0088; Wistar MIA = 5, IQR 1-6 versus WKY MIA = 15, IQR 8-28, p=0.0335). Although the response for the Aδ fibre latency tended to be higher in MIA-treated rats when compared to their within-strain saline-treated controls, no statistically significant differences were observed. A trend towards increased responses in the C-fibre latency range was present in WKY rats when compared to Wistar rats (**C**), with the largest number of responses recorded in MIA-treated WKY rats (58, IQR 21-64), no statistically significant differences were evident. Data represent the average number of action potentials recorded within each post-stimulus time frame, with individual data points shown, and bars representing median values and error bars the IQR.

### p<0.01 versus Wistar saline, + p<0.05 versus Wistar MIA, Kruskal-Wallis test with Dunn's multiple comparisons *post-hoc* testing.
